# A novel small molecule LLL12B inhibits STAT3 signaling and sensitizes ovarian cancer cell to paclitaxel and cisplatin

**DOI:** 10.1101/2020.09.22.307645

**Authors:** Ruijie Zhang, Xiaozhi Yang, Dana M. Roque, Chenglong Li, Jiayuh Lin

**Affiliations:** Department of Thoracic Surgery, Tongji Hospital, Tongji Medical College, Huazhong University of Science and Technology, Wuhan, P.R. China; Department of Biochemistry and Molecular Biology, University of Maryland School of Medicine, Baltimore, MD, USA; Department of Medicinal Chemistry, College of Pharmacy, the University of Florida, Gainesville, FL 32610, USA; Division of Gynecologic Oncology, Department of Obstetrics and Gynecology, University of Maryland School of Medicine, Baltimore, MD, USA

**Keywords:** Small molecule inhibitors, STAT3, ovarian cancer, paclitaxel, cisplatin

## Abstract

Ovarian cancer is the fifth most common cause of cancer deaths among American women. Platinum and taxane combination chemotherapy represents the first-line approach for ovarian cancer, but treatment success is often limited by chemoresistance. Therefore, it is necessary to find new drugs to sensitize ovarian cancer cells to chemotherapy. Persistent activation of Signal Transducer and Activator of Transcription 3 (STAT3) signaling plays an important role in oncogenesis. Using a novel approach called advanced multiple ligand simultaneous docking (AMLSD), we developed a novel nonpeptide small molecule, LLL12B, which targets the STAT3 pathway. In this study, LLL12B inhibited STAT3 phosphorylation (tyrosine 705) and the expression of its downstream targets, which are associated with cancer cell proliferation and survival. We showed that LLL12B also inhibits cell viability, migration, and proliferation in human ovarian cancer cells. LLL12B combined with either paclitaxel or with cisplatin demonstrated synergistic inhibitory effects relative to monotherapy in inhibiting cell viability and LLL12B-paclitaxel or LLL12B-cisplatin combination exhibited greater inhibitory effects than cisplatin-paclitaxel combination in ovarian cancer cells. Furthermore, LLL12B-paclitaxel or LLL12B-cisplatin combination showed more significant in inhibiting cell migration and growth than monotherapy in ovarian cancer cells. In summary, our results support the novel small molecule LLL12B as a potent STAT3 inhibitor in human ovarian cancer cellsand suggest that LLL12B in combination with the current front-line chemotherapeutic drugs cisplatin and paclitaxel may represent a promising approach for ovarian cancer therapy.

## Introduction

Ovarian cancer is the most lethal gynecologic malignancy [1-2]. In 2018, there were approximately 22,240 new cases and 14,070 deaths from ovarian cancer in the United States [3]. Because of a lack of early symptoms, nearly 80% of patients will be diagnosed at an advanced stage [4]. Adjuvant chemotherapy is usually needed following surgical cytoreduction. Platinum in combination with taxane chemotherapy is considered a first-line approach [5-7]. Unfortunately, five-year survival rates have not improved much during the past 20 years [8-11], due to both intrinsic and acquired chemoresistance. Many research efforts focus upon reversal of chemotherapy resistance for recurrent disease, but less attention is given to enhancing sensitivity to chemotherapy during the primary treatment of ovarian cancer [12-15]. In recent years, many researchers have invested in biotherapies with more precise targets, including immunotherapy, gene therapy, and molecular targeted therapy, which may be more effective and less toxic [16]. As the most common member of the signal transducers and activators of transcription (STAT) proteins, STAT3 is critical to many signaling pathways [17]. In normal cells, the activation of STAT3 is transitory and restricted; it can promote embryo development and growth, induce inflammation, and cause autophagy, among other processes [18]. STAT3 also plays an important role in the development of tumors, and is now considered an oncogene. The constitutive and abnormal activation of STAT3 can upregulate or downregulate many target tumor-related genes, such as *BCL-2, c-myc, cyclinD1, survivin, cleaved caspase-3, HIF-1* and *VEGF*, which then enable various processes key to malignant progression, such as cell proliferation, tumor initiation, migration, invasion, angiogenesis, metastasis, cell cycle dysregulation, induction of the epithelial mesenchymal transition (EMT), and inhibition of apoptosis, as well as promote multidrug resistance to chemotherapy [18]. STAT3 activation occurs when the tyrosine 705 (Tyr705) residue is phosphorylated. Using a novel approach called advanced multiple ligand simultaneous docking (AMLSD), we developed several new small molecular inhibitors targeting STAT3, including the novel STAT3 inhibitor LLL12B. Computer models with docking simulation showed that LLL12B binds directly to the phosphoryl tyrosine 705(pTyr705) binding site of the STAT3 monomer. In the present study, we characterized the biologic effects of LLL12B alone and in combination with chemotherapy on ovarian cancer cells.

## Material and Methods

### Materials

The small molecule LLL12B, a novel STAT3 inhibitor, was synthesized at University of Florida College of Pharmacy (Chenglong Li). LLL12B powder was dissolved in sterile dimethyl sulfoxide (DMSO) to make a 20 mM stock solution and stored at -20 °C. Cisplatin and 3-(4, 5- dimethylthiazol-2-yl)-2, 5-diphenyltetrazolium bromide (MTT) were purchased from Sigma (Burlington, MA). The stock concentration of cisplatin was 5mM in ddH2O. Paclitaxel was obtained from LC Laboratories (Woburn, MA). The stock concentration of paclitaxel was 20mM in DMSO. Primary and secondary antibodies were bought from Cell Signaling Technology (Danvers, MA).

### Cell Lines

All four human ovarian cancer cell lines (A2780, SKOV3, CAOV3, and OVCAR5), were purchased from ATCC (American Type Culture Collection, Manassas, VA). OVAR5 was cultured in RPMI1640 medium with 10% fetal bovine serum (FBS) and 1% penicillin/streptomycin (PS); A2780, CAOV3, and SKOV3 were cultured in Dulbecco’s modified Eagle medium (DMEM) with 10% FBS and 1% PS. All of the cell lines were maintained in a humidified 37°C incubator with 5% CO_2_/95% air. Media were replaced twice a week.

### Western blot analysis

The four cell lines were seeded in 10cm-plates in 70% cell density, then treated with DMSO or different concentrations of LLL12B. Cells were cultured overnight before they were harvested for Western blot analysis. Ovarian cancer cells were lysed in cold lysis buffer and the proteins were separated by 10% SDS-PAGE. Proteins were transferred to PVDF membrane under 350mA for 110 minutes and then blocked by 5% milk for 1 hour and incubated overnight at 4°C with antibodies: anti p-STAT3 (Tyrosine 705), GAPDH, c-MYC, cylinD1, survivin, and cleaved caspase 3. After washing with tris-buffered saline-tween (TBST) for 3 times (15 min), the membranes were blotted with the secondary antibody and scanned with a Storm Scanner (Amersham Pharmacia Biotech Inc, Piscataway, NJ).

### MTT cell viability assay

We seeded A2780, SKOV3, CAOV3 and OVCAR5 in 96-well microtiter plates at a density of 3,000 cells with 100μl medium per well. After overnight incubation, the cells were treated in each well at 37°C with vehicle control (DMSO) or different concentrations of drugs: LLL12B, cisplatin alone, paclitaxel alone, or their combination. Seventy-two hours later, we added 20μl of MTT to each well. After incubation for 4 h at 37°C, each well was supplemented with 150μl of dimethylformamide solubilization solution followed by an incubation overnight, protected from light at room temperature. Cell viability was assessed using the absorbance at 595 nm of each well. The DMSO cells were set at 100% and the cell viability of drug-treated cells was determined relative to DMSO cells. Then the combination index (CI) was determined by CompuSyn software (www.combosyn.com). The CI values indicate an additive effect when equal to 1, an antagonistic effect when >1, and a synergistic effect when <1 based on the theorem of Chou and Talalay [19].

### Wound-healing/cell migration assay

A2780 and SKOV3 cells were seeded in 6-well plates and incubated at 37°C overnight. When cells reached 100% confluence, the monolayer was scratched by a 100-μl pipette tip. We washed each well with PBS twice and added new medium with different drugs: DMSO, LLL12B, paclitaxel, cisplatin or their combination. Photos of each well were captured by microscope at time zero. Cells were incubated at 37°C until the wound of the control well was healed (SKOV3, 18h; A2780, 56h). The photos of each well were captured by microscope again after washing in PBS twice. Inhibition of migration was measured by ImageJ software (http://rsb.info.nih.gov/ij/) and calculated by the formula: percent of wound healed = 100 − [(final area / initial area) × 100%] [20].

### Cell growth assay

All four cell lines were seeded in 12-well plates at the same cell density, which was dependent on the growth ability of each cell line (SKOV3:1*10^4^ cells per well, CAOV3: 2.5*10^4^ cells per well, A2780:0.5*10^4^ cells per well, and OVCAR5: 1*10^4^ cells per well). The cells were cultured overnight at 37°C, then were treated with DMSO or different concentrations of drugs: LLL12B, paclitaxel alone, cisplatin alone, or the combination. We counted the cell number of each well days 2, 4, and 6 after treatment to generate growth curves. Significant differences were defined as **p<0.01 and ****p<0.0001.

### Statistical analysis

Significance of correlations was determined by GraphPad Prism 7 software (GraphPad Software Inc., San Diego, CA, USA). The data were expressed as mean ± standard deviations (SD). One-way ANOVA and two-way ANOVA with Tukey’s Test were used to analyze the statistical difference between two groups. Significance was set at *p* < 0.05. The *, ** and *** indicate *p* < 0.05, *p* < 0.01 and *p* < 0.001, respectively.

## Results

### LLL12B inhibited STAT3 phosphorylation and the expression of downstream targeted genes

To target STAT3, we used a novel approach called advanced multiple ligand simultaneous docking (AMLSD), and developed a novel nonpeptide small molecule, LLL12B. The chemical structure of LLL12B is shown (Figure 1). In order to examine the ability of LLL12B to inhibit p-STAT3 (Tyr705) *in vitro*,Western blotting analysis was performed by us. All four ovarian cancer cell lines were seeded in 10 cm plates and were treated with DMSO or different concentrations of LLL12B (0.25uM-2.5uM). Protein expression levels were analyzed. Compared with those treated by DMSO, p-STAT3 was inhibited by LLL12B (Figure 2). In addition, the STAT3 downstream targets c-myc, cyclinD1, and survivin were down-regulated and cleaved caspase-3 was induced (Figure 1). LLL12B inhibited STAT3 phosphorylation and down-regulated downstream target genes which are associated with cancer cell proliferation and growth [21-24]. These results indicate that LLL12B is a biologically relevant potent STAT3 inhibitor of ovarian cancer cells.

**Figure 1.**
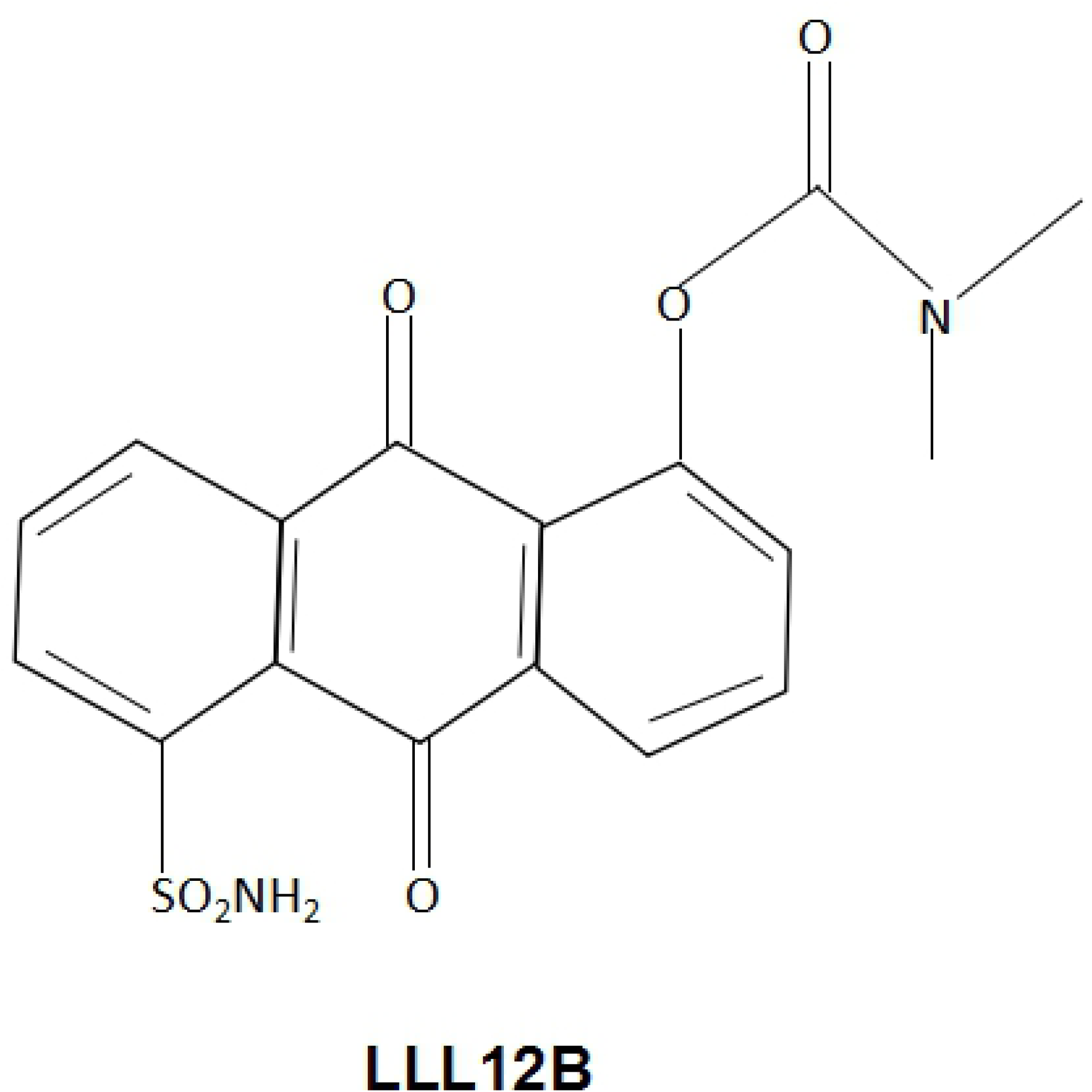
The chemical structure of LLL12B.

**Figure 2.**
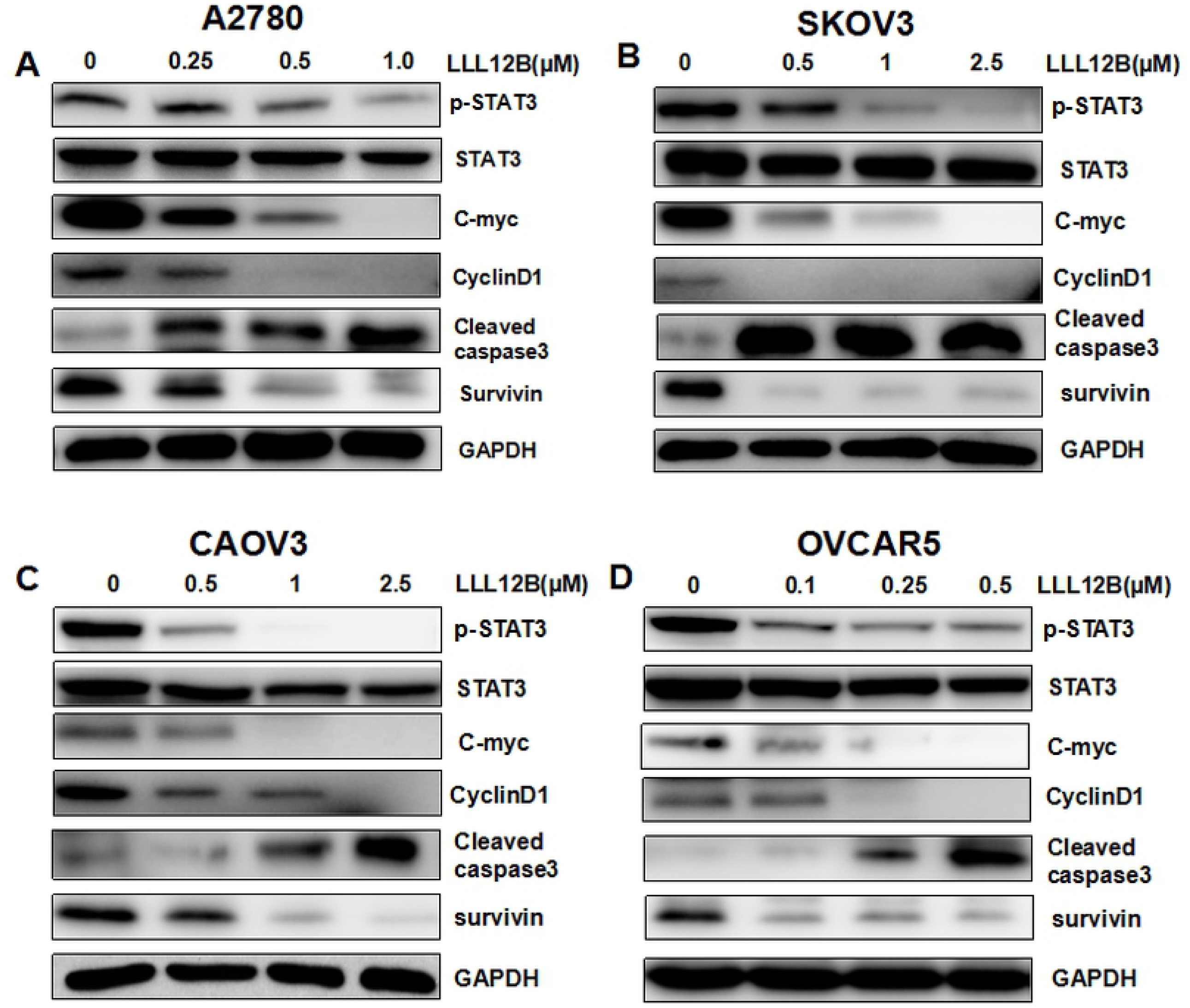
LLL12B inhibits p-STAT3 and its downstream targets in human ovarian cancer cells. **A-D** The effects of LLL12B on STAT3 phosphorylation (Tyr705) and its downstream targets in ovarian cancer cell lines. A2780, CAOV3, SKOV3, OVCAR5 cells were treated with DMSO or different concentrations of LLL12B. The levels of p-STAT3 and the downstream target gene proteins were determined by Western blots.

### LLL12B inhibited cell viability of human ovarian cancer cells and synergistically enhanced the effect of cisplatin and paclitaxel

To evaluate inhibition of cell viability, MTT assays were performed using A2780, CAOV3, SKOV3 and OVCAR5 cells. Cells were seeded in 96-well plates and treated with LLL12B or DMSO control followed by culture at 37°C for additional 72 hours. Cell viability was significantly inhibited by LLL12B (Figure 3). To investigate whether chemotherapy can be enhanced by LLL12B, the cells were treated by LLL12B combined with cisplatin or paclitaxel. The combination index (CI) showed that the suppression achieved with combination treatment was more significant than that of any monotherapy. The CIs of LLL12B combined with cisplatin or paclitaxel in each cell line were all less than 1, which indicated synergism. Furthermore, the combination of LLL12B with cisplatin or paclitaxel exhibited more significant inhibitory effects on cell viability than the combination of cisplatin and paclitaxel (Figure 3). These results indicate that LLL12B inhibits cell viability and also synergistically enhances the effect of chemotherapy in ovarian cancer cell lines.

**Figure 3.**
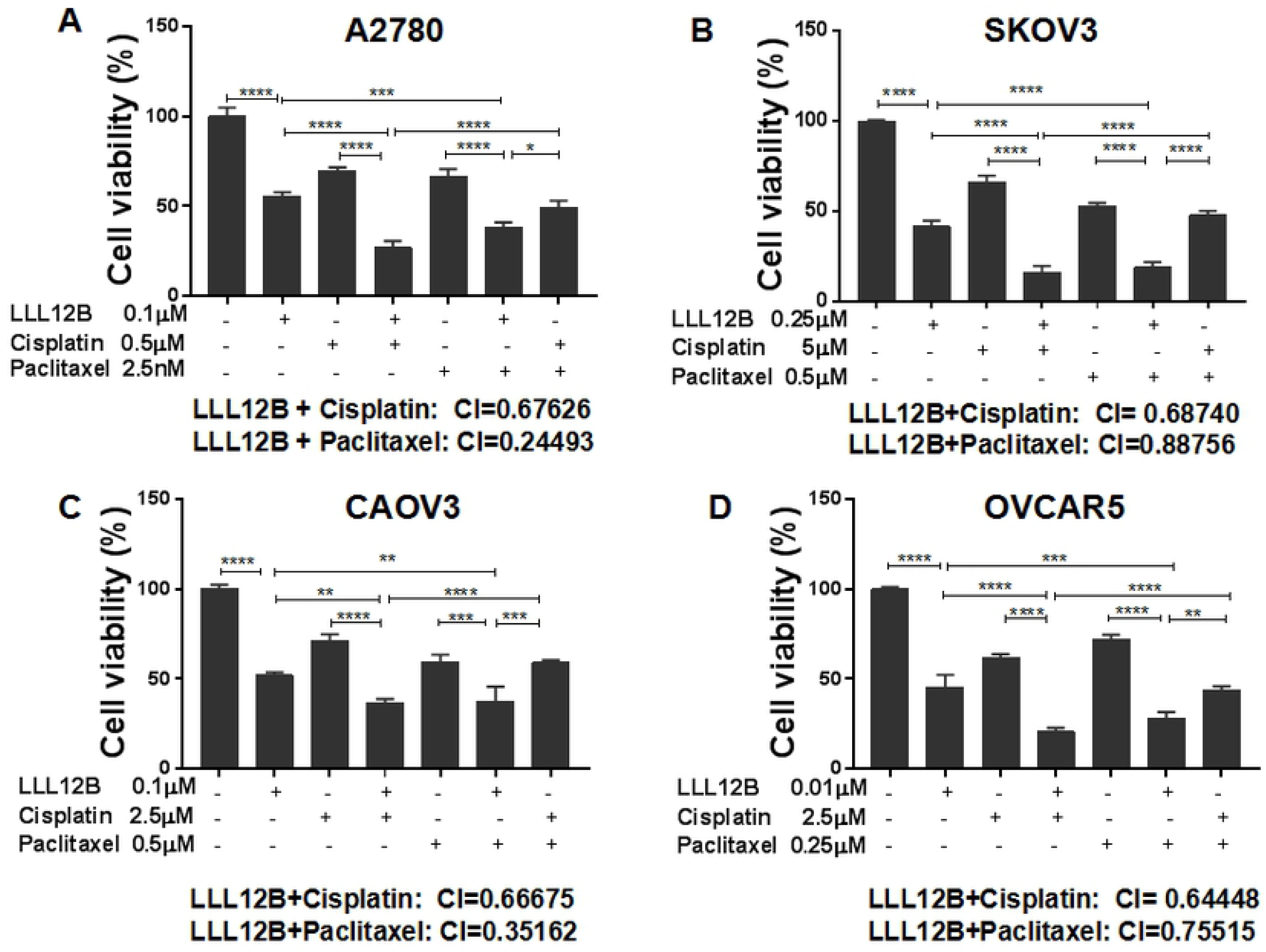
The effects of LLL12B, cisplatin, paclitaxel, and drug combination on cell viability. MTT assays were performed to evaluate cell viability. **A-D** LLL12B inhibited cell viability of ovarian cancer cells, which were synergistically inhibited when LLL12B was combined with cisplatin or paclitaxel. The differences were found to be significantly different at *p<0.05, **p<0.01,***p<0.001 and ****p<0.0001.

### LLL12B inhibited cell migration of ovarian cancer cells and enhanced the effect of cisplatin and paclitaxel

Cell migration is an important step in tumor invasion and metastasis, which confers prognosis. According to previous literature reports [43], cancer cell migration can be inhibited by blocking of STAT3 pathway. Therefore, we tested the effects of the novel STAT3 inhibitor LLL12B on ovarian cancer cell migration and whether the effect of cisplatin and paclitaxel can be enhanced by LLL12B. Only A2780 and SKOV3 cell lines were tested because the monolayer phenotypes of the other two cell lines were not suitable for cell migration assays. Compared with the DMSO control, cell migration was inhibited by LLL12B. The combination of LLL12B with cisplatin or paclitaxel resulted in more significant inhibition of cell migration compared to monotherapy; notably, the inhibitory effects exceeded that of paclitaxel and cisplatin in combination (Figure 4). These results indicate that LLL12B can inhibit cell migration of ovarian cancer cells, and may also enhance the effects of chemotherapy. This suggests that LLL12B may be helpful for treatment or prevention of ovarian cancer cell invasion and metastasis.

**Figure 4.**
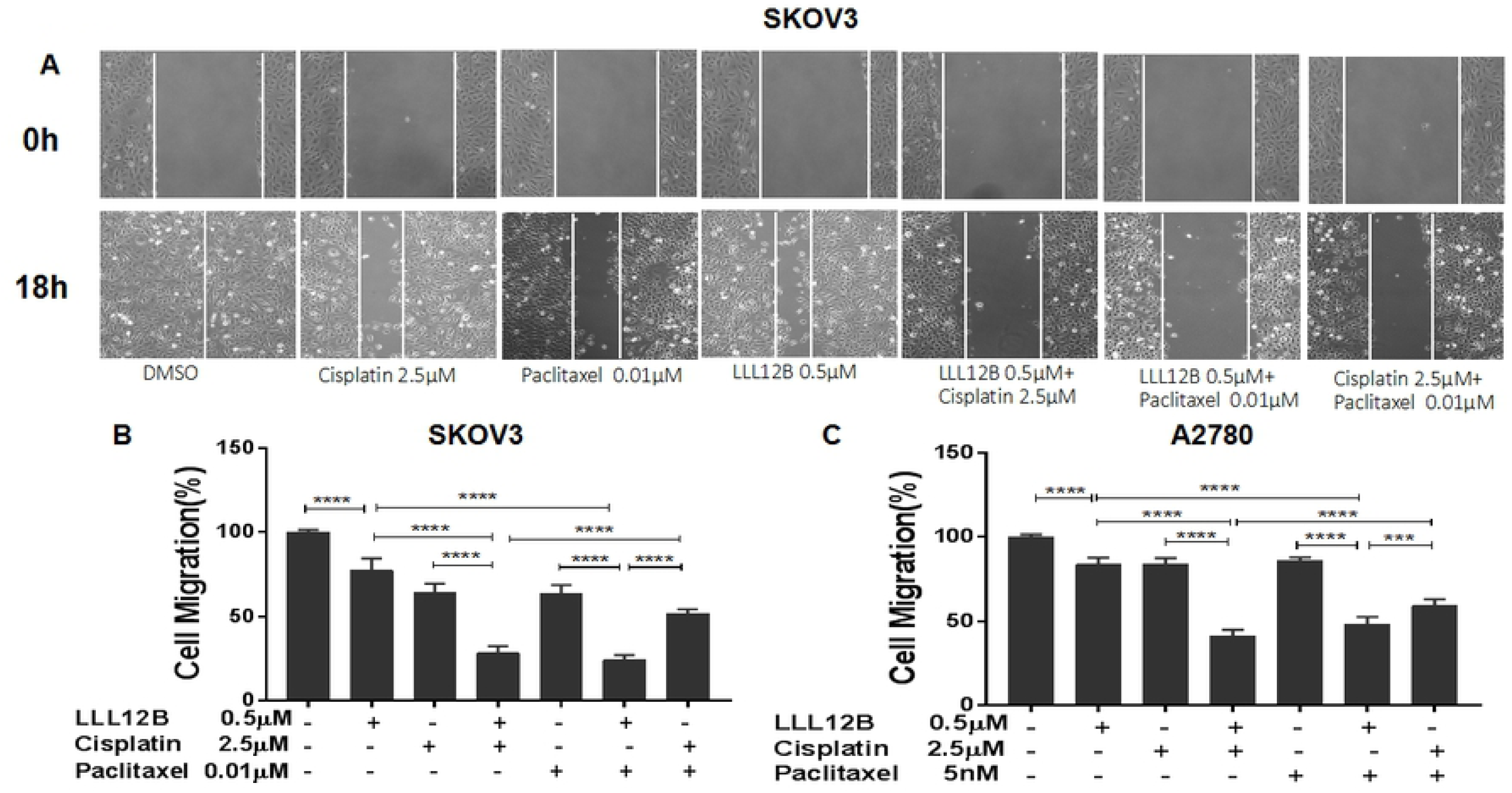
The effects of LLL12B, cisplatin, paclitaxel, and drug combination on cell migration. Wound-healing assays were performed to evaluate the migration ability of SKOV3 and A2780 ovarian cancer cells. SKOV3 and A2780 cells were seeded in 6-well plates and treated with DMSO or different concentration of drugs: LLL12B or cisplatin or paclitaxel or the combination. The differences were found to be significantly different at ***p<0.001 and ****p<0.0001.

### LLL12B inhibited cell growth and enhanced the effect of cisplatin and paclitaxel

Since LLL12B synergistically inhibited cell viability of ovarian cancer cells treated with cisplatin and paclitaxel, we then sought to investigate whether LLL12B could also inhibit cell growth using standard growth curves. Our results showed that LLL12B inhibited cell growth in all four ovarian cancer lines (Figure 5,Figure 6). Parallel to our observations for viability, combination treatment of LLL12B with cisplatin or paclitaxel resulted in greater inhibitory effects on cell growth than monotherapy (Figure 5,Figure 6).

**Figure 5.**
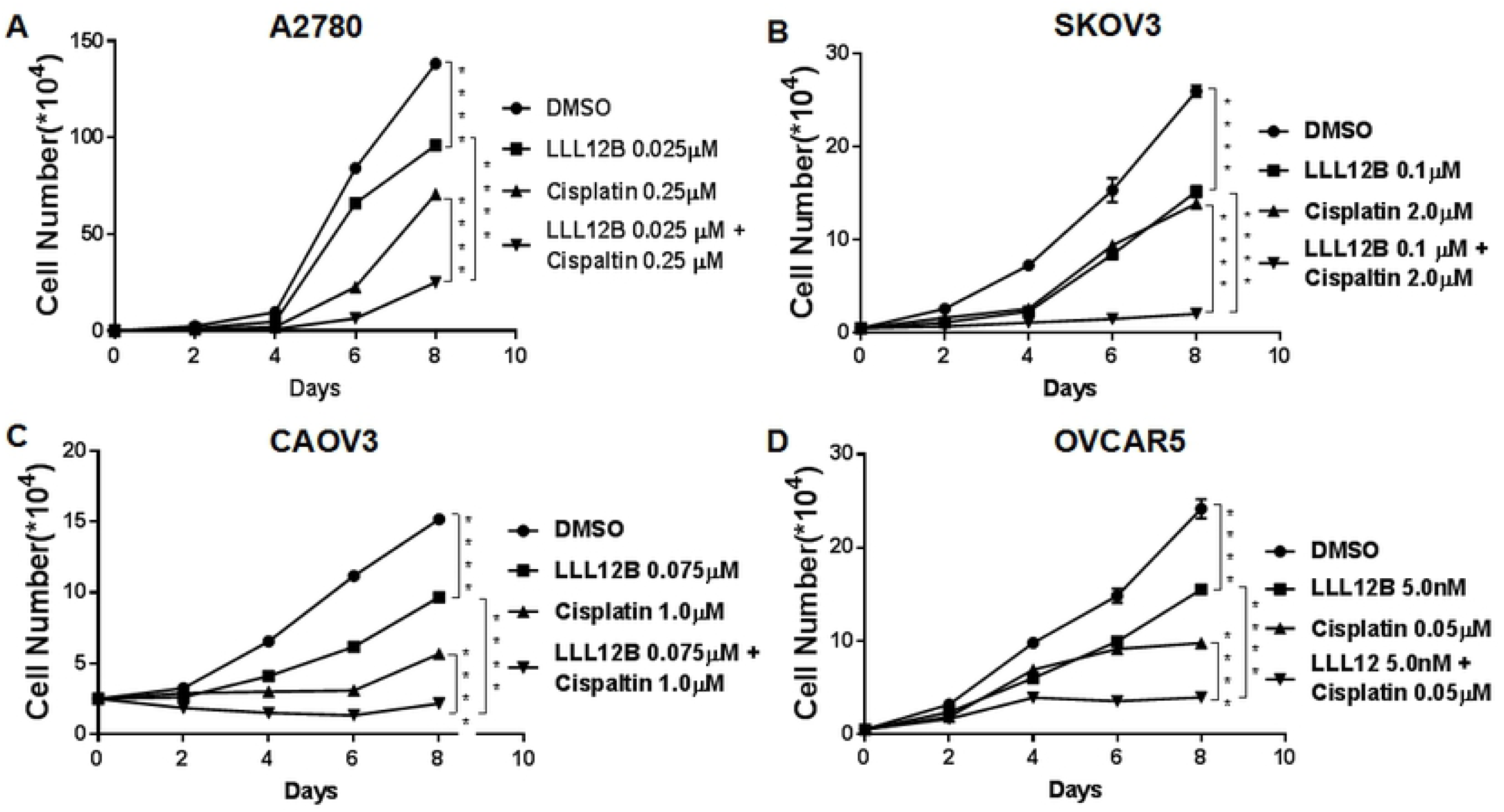
The effects of LLL12B, cisplatin, and drug combination on cancer cell growth. Cell growth assays were performed to evaluate cell proliferation ability of ovarian cancer cells. Cells were treated with LLL12B, cisplatin and their combination. The differences were found to be significantly different at **p<0.01 and ****p<0.0001. LLL12B alone or combined with cisplatin inhibited cell growth of ovarian cancer cells.

**Figure 6.**
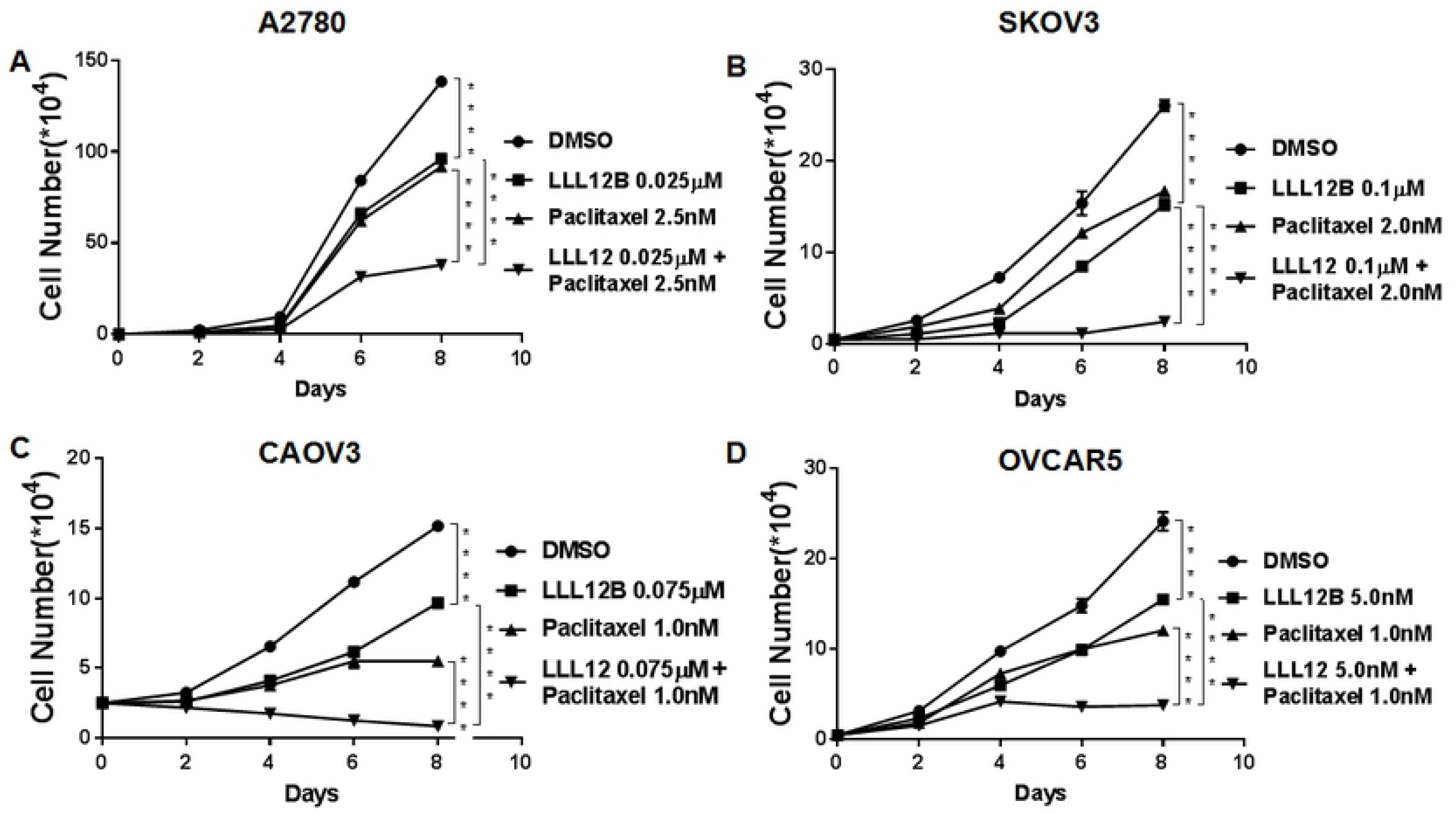
The effects of LLL12B, paclitaxel, and drug combination on cancer cell growth. Cell growth assays were performed to evaluate cell proliferation ability of ovarian cancer cells. Cells were treated with LLL12B, paclitaxel and their combination. The differences were found to be significantly different at **p<0.01 and ****p<0.0001.LLL12B alone or combined with paclitaxel inhibited cell growth of ovarian cancer cells.

## Discussion

Ovarian cancer is the tenth most common cancer and the fifth most lethal cancer of women in the United States [1-2]. Among gynecologic malignancies, ovarian cancer has the highest mortality rate. Ovarian cancer is very difficult to diagnose early, and more than 50% of patients are diagnosed with advanced (stage III/IV) disease [1-3]. The standard treatment for advanced ovarian cancer is radical resection combined with platinum-taxane combination chemotherapy [4, 26-27]. Despite small improvements in 5-year overall survival for this disease, there are still many challenges for its treatment. The 5-year overall survival of advanced-stage ovarian cancer is no more than 20–35%, and has not changed much for nearly twenty years. Both intrinsic and acquired chemoresistance remain problematic, and as many as 75% of ovarian cancer patients suffer from cancer recurrence [3]. It is therefore crucial to find new drugs to enhance the effect of current chemotherapy treatment. Targeted therapy often offers the benefit of precise action and fewer side effects.

As an important transducer of many cytokines and growth factors, STAT proteins have 7 members (STAT1, STAT2, STAT3, STAT4, STAT5A, STAT5B and STAT6), and STAT3 is the most widely known and researched [28]. Compared with normal tissues, STAT3 is overexpressed or constitutively activated in about 70% of human solid tumors and in 94% of ovarian cancers [19,29-31].. A homodimer is formed when STAT3 is activated by phosphorylation.,which translocates to the nucleus,recognizes and binds to STAT3-specific DNA-binding elements.Then some target genes can be regulated to promote cell growth,prevent apoptosis and so on.[17,32-34]. Abnormal activation of STAT3 can induce malignant cell transformation and is related to the poor prognosis of certain tumors. Conversely, the disruption of constitutively activated STAT3 can promote cell apoptosis and suppress tumor-cell growth.

At the same time, it is reported that over expression of STAT3 also associated with cisplatin resistance and paclitaxel resistance [17,35-37]. We have previously demonstrated that constitutive activation of STAT3 was present in ovarian cancer cell lines but not in normal ovarian surface epithelial cells [38], making selective STAT3 inhibition an excellent candidate for ovarian cancer treatment. In this study, we provide evidence that STAT3 inhibition may be a good enhancer for cisplatin and paclitaxel chemotherapy.

The SH2 domain is a critical module among the six structural domains of STAT3, which facilitate binding to specific p-Tyrosine (Tyr) motifs of receptors for activation of the protein. Interaction of the pTyr-SH2 domain with STAT3 dimerization represents an important molecular event for STAT3 functioning. For these reasons, most drugs have been designed to bind this domain. Many peptide-based inhibitors of STAT3 have been reported, but their use is limited by poor cell permeability and limited *in vivo* stability [39-40]. During recent years, many nonpeptide small molecule inhibitors have been developed, which show better stability [41-42]. We previously developed several nonpeptide small molecule STAT3 inhibitors, such as LLL12, which inhibits STAT3 phosphorylation and suppresses the development of cancer [43-46]. In the present study, we explored LLL12B, a carbamate-based prodrug for LLL12. LLL12B has one of the smallest molecular weights (374 dalton) compared to other STAT3 inhibitors. In addition, our *in vivo* pharmacokinetic studies in rats (data not shown) indicated that LLL12B is orally bioavailable (38.0%) and stable in the plasma, producing drug levels 5-fold better compared to LLL12. These results support that LLL12B is a superior drug relative to LLL12 to target STAT3 in ovarian cancer cells.

In this study, we tested LLL12B in several well-characterized human ovarian cancer cell lines. LLL12B consistently inhibited STAT3 phosphorylation and downregulated the downstream targets. LLL12B exerted potent inhibition of cell viability, migration and growth. When cisplatin or paclitaxel was combined with LLL12B, inhibition of these parameters was enhanced relative to monotherapy and, importantly, greater than that of paclitaxel with cisplatin, which currently represents the standard of care.

In conclusion, the novel small molecule STAT3 inhibitor, LLL12B, designed by AMLSD methodology shows excellent therapeutic potential in ovarian cancer cell lines. Our results suggest that LLL12B is a potent STAT3 inhibitor in ovarian cancer, and that LLL12B in combination with the current front-line chemotherapeutic drugs cisplatin and paclitaxel may represent a promising approach for ovarian cancer therapy that warrants further study.

## Acknowledgements

We thank Dr. Richard Eckert at the Department of Biochemistry and Molecular Biology at the University of Maryland (Baltimore, MD, USA) for providing the microscope used to evaluate the wound healing assay.

## Funding

This research was supported by the University of Maryland School of Medicine and Greenebaum Comprehensive Cancer Center start up fund.

## Competing interests

The authors declare no conflict of interest.

